# A CRISPR-Cas9-mediated F0 screen to identify pro-regenerative genes in the zebrafish retinal pigment epithelium

**DOI:** 10.1101/2022.08.28.505611

**Authors:** Fangfang Lu, Lyndsay L. Leach, Jeffrey M. Gross

**Affiliations:** Department of Ophthalmology, Louis J. Fox Center for Vision Restoration, University of Pittsburgh School of Medicine, Pittsburgh PA, 15213; Department of Developmental Biology, University of Pittsburgh School of Medicine, Pittsburgh PA, 15213; Department of Ophthalmology, The Second Xiangya Hospital, Central South University, Changsha, Hunan 410011 China; Department of Molecular Biosciences, The University of Texas at Austin, Austin TX 78712

**Keywords:** Retinal pigment epithelium, zebrafish, regeneration, CRISPR-Cas9, F0 screen, claudin-7

## Abstract

Ocular diseases resulting in death of the retinal pigment epithelium (RPE) lead to vision loss and blindness. There are currently no FDA-approved strategies to restore damaged RPE cells. Stimulating intrinsic regenerative responses within damaged tissues has gained traction as a possible mechanism for tissue repair. Zebrafish possess remarkable regenerative abilities, including within the RPE; however, our understanding of the underlying mechanisms remains limited. Here, we conducted an F0 *in vivo* CRISPR-Cas9-mediated screen of 27 candidate RPE regeneration genes. The screen involved injection of a ribonucleoprotein complex containing three highly mutagenic guide RNAs per target gene followed by PCR-based genotyping to identify large intragenic deletions and MATLAB-based automated quantification of RPE regeneration. Through this F0 screening pipeline, eight positive and seven negative regulators of RPE regeneration were identified. Further characterization of one candidate, *cldn7b*, revealed novel roles in regulating macrophage/microglia infiltration after RPE injury and in clearing RPE/pigment debris during late-phase RPE regeneration. Taken together, these data support the utility of targeted F0 screens for validating pro-regenerative factors and reveal novel factors that could regulate regenerative responses within the zebrafish RPE.

**Summary statement:** We present a rapid CRISPR-Cas9-mediated F0 screen, which revealed novel regulators of retinal pigment epithelium (RPE) regeneration in zebrafish. This screen and identified factors will greatly facilitate discovery of underlying RPE regenerative mechanisms.

## INTRODUCTION

The retinal pigment epithelium (RPE) is a monolayer of pigmented cells that function to maintain the health of the neural retina and support vision (Lakkaraju *et al*., 2020). Dysfunction of the RPE contributes to many ocular degenerative diseases, including age-related macular degeneration (Sparrow *et al*., 2010). Mammals have a limited ability to repair RPE damage and therefore RPE dysfunction inevitably results in permanent visual impairment (Grierson *et al*., 1994; Del Priore *et al*., 1995). An intriguing therapeutic approach to overcome irreversible RPE damage is to leverage the intrinsic proliferative, and possibly regenerative, potential of RPE *in situ* (Longbottom *et al*., 2009; Xia *et al*., 2011; Salero *et al*., 2012; Chen *et al*., 2016; Kampik *et al*., 2017). For this to become a reality, we must understand the mechanisms driving intrinsic RPE regeneration. Zebrafish possess robust regenerative competence and share genetic synteny with humans (Howe *et al*., 2013). Thus, knowledge gained from identifying mechanisms driving RPE regeneration in zebrafish can be applied to non-regenerative mammals and, eventually, to humans. Previous work from our laboratory demonstrated the capacity for intrinsic RPE regeneration in zebrafish after widespread genetic ablation (Hanovice *et al*., 2019). Subsequent RNA-sequencing (RNA-seq) experiments identified an extensive list of potential regulators of RPE regeneration, including mTOR (Lu *et al*., 2022) and immune (Leach *et al*., 2021) pathway responses. Many genes remain to be investigated *in vivo* and high-throughput methodology is needed to screen these candidates.

Among the existing genome editing tools, <u>c</u>lustered regularly interspaced <u>s</u>hort <u>p</u>alindromic repeats (CRISPR)-Cas9 technology has emerged as a facile system for generating mutations in whole organisms (Hwang *et al*., 2013; Hsu *et al*., 2014; Zuo *et al*., 2017; Liu *et al*., 2019). This utility is due to the simplicity of designing single guide RNAs (sgRNAs), the efficiency and precision of targeting virtually any genomic locus, and the ease of multiplex genome editing, among other features (Ran *et al*., 2013; Unal Eroglu *et al*., 2018). As a result, CRISPR-Cas9 technology has been widely applied in zebrafish to systematically investigate gene function (Varshney *et al*., 2015; Pei *et al*., 2018). Historically, however, phenotypes have often been analyzed in the F2 generation, when homozygous mutants are obtained (Varshney *et al*., 2016; Sorlien *et al*., 2018), extending time to discovery and limiting throughput. F0 screens have emerged as a rapid method for identifying phenotypes, but the efficacy of these screens depends on the mutagenesis efficiency of CRISPR-Cas9. Recently, several advancements have improved targeted F0 mutagenesis, including multiplexed pooled injections of gRNAs (Shah *et al*., 2015), use of pre-assembled Cas9-sgRNA ribonucleoprotein complexes (RNPs) (Burger *et al*., 2016; Kuil *et al*., 2019), use of synthetic two-RNA components (crRNA:tracrRNA) guide RNAs (gRNAs) (Hoshijima *et al*., 2019), and multi-locus targeting (Wu *et al*., 2018; Hoshijima *et al*., 2019; Keatinge *etal*., 2021; Kroll *et al*., 2021). Of interest here, the knockout methodology developed by Kroll *et al*. maximized CRISPR-Cas9 editing efficiency by combining multi-locus targeting with methodology to enhance mutagenesis at each locus (Kroll *et al*., 2021). Indeed, targeting three unique loci per gene with synthetic gRNAs consistently converted >90% of injected embryos directly into biallelic knockouts. Additionally, this study adapted a sequencing-free tool, headloop PCR (Rand *et al*., 2005), to rapidly validate the mutagenic activity of individual gRNAs. The high proportion of biallelic knockouts in the F0 populations coupled with economical PCR-based mutagenic screening make this an excellent approach for high-throughput screens.

Here, we selected 27 candidate RPE regeneration genes identified from previous RNA-seq studies (Leach *et al*., 2021; Lu *et al*., 2022) and combined our zebrafish RPE ablation-regeneration model (Hanovice *et al*., 2019) with CRISPR-Cas9 mediated F0 knockout technology to develop a scalable F0 screening pipeline to identify essential regulators of RPE regeneration. This pipeline is composed of the Kroll *et al*. F0 knockout methodology (Kroll *et al*., 2021), PCR amplification-based genotyping, and a MATLAB-based automated quantification platform, RpEGEN (Leach *et al*., 2022). Eight potential positive regulators and seven potential negative regulators of RPE regeneration were identified in the screen. Further characterization of one gene of interest (GOI), *cldn7b*, revealed an essential role in modulating the activity of macrophages/microglia during late-phase RPE regeneration.

## RESULTS AND DISCUSSION

### A CRISPR/Cas9-based F0 screening pipeline to identify genes that modulate RPE regeneration

Recent RNA-seq data from our laboratory have identified numerous factors with the potential to regulate RPE regeneration (Leach *et al*., 2021; Lu *et al*., 2022). Here, we sought to develop a pipeline to rapidly screen and validate gene candidates for roles in RPE regeneration. To achieve this, we selected 27 GOIs upregulated in RPE cells at 2 days post-injury (dpi) (Fig. S1) (Leach *et al*., 2021; Lu *et al*., 2022) based on predicted functions (e.g. cell-cell contact, lipid metabolism, immune responses) that are critical during tissue regeneration in a variety of organs/systems (Roy and Tedeschi, 2021; George *et al*., 2021). To generate knockouts, we modified an existing CRISPR/Cas9-based F0 mutagenesis methodology (Fig. 1A-D) (Kroll *et al*., 2021). For each GOI, three CRISPR RNAs (crRNAs) with Alt-R modifications were designed, each of which targeted non-overlapping regions of the 5’ exons (Fig. 1A). RNP complexes were preassembled *in vitro* using Cas9 protein and guide RNA (gRNA), which was composed of a crRNA and a trans-acting CRISPR RNA (tracrRNA) (Hoshijima *et al*., 2019). Headloop PCR (Rand *et al*., 2005) was used to pre-screen individual RNP complexes to ensure substantial mutagenesis at each locus (Fig. 1B,C). The mutagenic efficiency of each individual RNP was determined quantitatively, by assessing the ratio of headloop PCR bands (H), which indicate an effective mutagenesis event, to standard PCR bands (S); and qualitatively, by comparing H and S band intensities (Fig. 1C). gRNAs showing a mutagenic ratio >70% were used for phenotypic screening (Table S1).

**Fig. 1.**
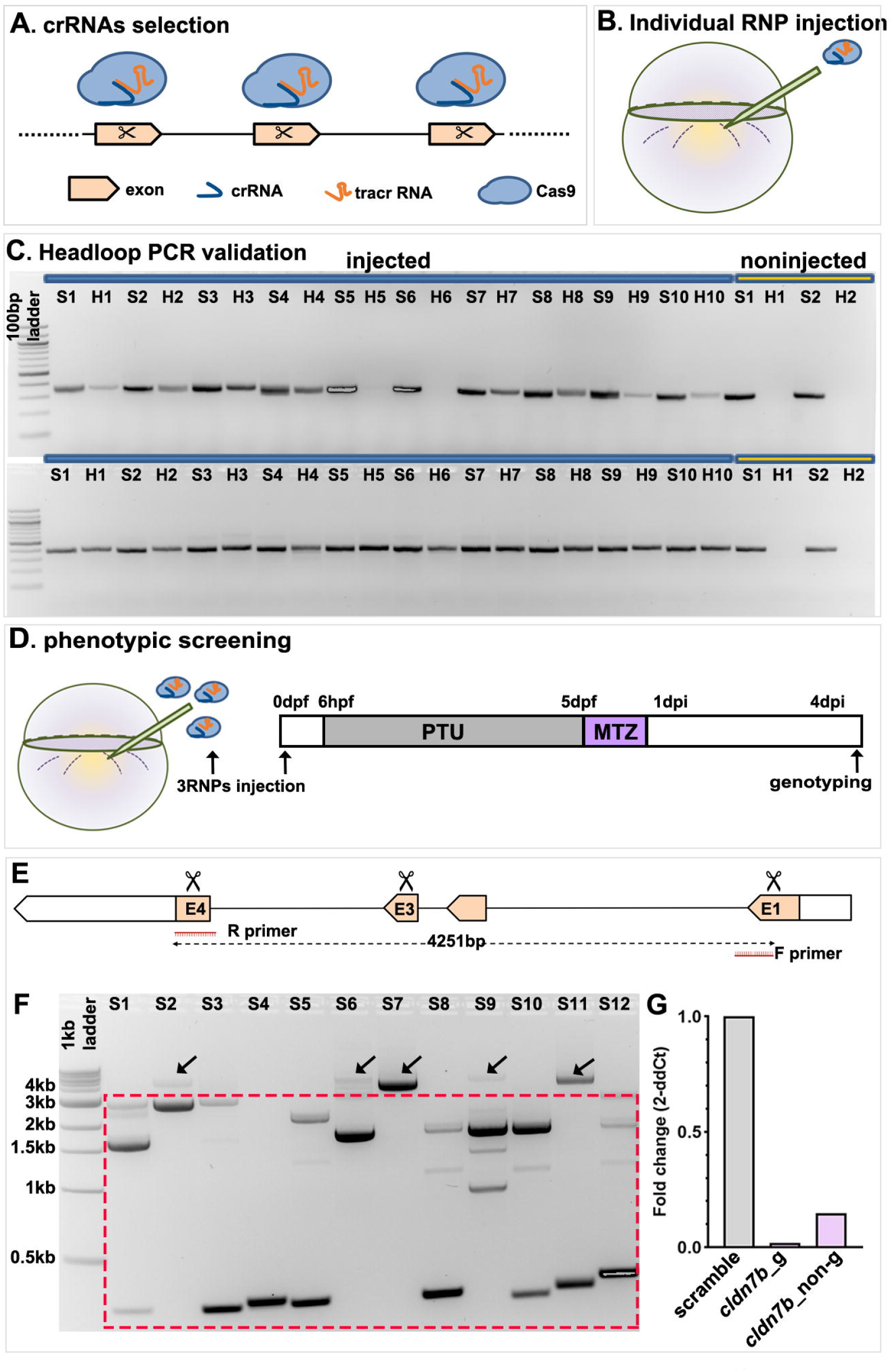
F0 screening pipeline to identify candidate genes that regulate RPE regeneration. (A) Design and selection of crRNAs targeting three non-overlapping regions of each candidate gene of interest (GOI). (B) Injection of individual gRNA (crRNA:tracrRNA) with Cas9 protein (RNP complex) into embryos derived from *rpe65a*:nfsB-eGFP transgenic crosses. (C) Example electrophoresis results of headloop PCR (H) targeting *zgc:153911* (top gel lanes) and *nrg1* (bottom gel lanes) in parallel with standard PCR (S) from ten RNP-injected embryos and two uninjected controls. The top gel lanes show a mutagenic rate of 80% for *zgc:153911*; the bottom gel lanes show a mutagenic rate of 100% for *nrg1*. Mutagenic rate = H+ /S+ number of embryos. (D) Schematic showing the MTZ-mediated RPE ablation timeline after injection of three RNPs for each GOI into one-cell stage embryos. (E) Schematic showing the sequence spanning all three target loci (scissors) in exon 1 (E1), exon 3 (E3), and exon 4 (E4) that would be PCR-amplified to confirm biallelic mutations. (F) A typical electrophoresis result (*cldn7b*) from twelve pooled RNP-injected larvae shows the large intragenic deletions generated by simultaneously targeting three genomic sites. Gel bands within the dashed box indicate the intragenic deletions. The bands above the box with a size of around 4kb (indicated by arrows) represent either wild-type alleles or mutated alleles with small indels that cannot be distinguished on the gel. (G) Bar graph showing relative expression fold change of *cldn7b* gene from genotyped (*cldn7b*_g), non-genotyped (*cldn7b*_non-g) *cldn7b* F0 knockout larvae and scrambled RNP-injected larvae. Each bar represents biological variation from n=50 pooled injected larvae and technical variation from three wells. Abbreviations as follows: RNP, ribonucleoprotein; S, standard PCR; H, headloop PCR; PTU, n-phenylthiourea; MTZ, metronidazole; hpf/dpf, hour/day (s) post-fertilization; dpi, day (s) post-injury; E, exon; F primer, forward primer; R primer, reverse primer.

After confirming effective mutagenesis of individual RNPs, a pool of three RNPs per GOI was injected into one-cell stage embryos, while control embryos were injected with a pool of three scrambled RNPs that were predicted to not target the zebrafish genome (Fig. 1D). Regardless of the gene targeted, pooled RNP injections resulted in dead and/or dysmorphic embryos, with incidences ranging from 13.3% to 40.5% (Table S1), consistent with published results (Kroll *et al*., 2021). To confirm biallelic mutations in F0 knockout larvae used for phenotypic analysis, we performed PCR amplification of sequences spanning all three target sites in the GOI, with the expectation that large intragenic deletions caused by the joining of sequences distal to the single RNP-induced double-strand breaks would be detectable (Fig. 1E). Indeed, large deletions were detected in most injected embryos (Fig. 1F). These are likely to represent loss-of-function mutations that prevent the expression of the protein of interest (Hoshijima *et al*., 2019; Kim and Zhang, 2020) and/or functional rescue through genetic compensation (El-Brolosy *et al*., 2019). For 20 of the 27 GOIs, only genotyped F0 larvae carrying large intragenic deletions were used for subsequent phenotypic analyses. For the remaining 7 GOIs, it was not feasible to PCR amplify across target sites due to the size of the locus (Table S1). Therefore, while the individual RNPs were each validated for these 7 GOIs, the combined efficiency of the three pooled RNPs was not assessed. To further validate our F0 knockout pipeline, we used quantitative real-time PCR (qRT-PCR) to measure mRNA levels of one GOI, *cldn7b*, in genotyped F0 knockout larvae. As anticipated, qRT-PCR results showed an approximately 98% decrease in *cldn7b* expression in genotyped *cldn7b* F0 knockout larvae with large deletions, when compared to scrambled controls (Fig. 1G). Moreover, pooled/non-genotyped *cldn7b* F0 knockout larvae showed an approximately 85% loss of *cldn7b* expression when compared to scrambled controls, suggesting that the pooled RNPs were highly effective in mutagenizing the target locus and reducing gene expression, even when large intragenic deletions were not pre-screened and enriched. Taken together, these results establish an efficient and robust F0 screening pipeline for use in identifying factors that contribute to RPE regeneration.

### Phenotypic screening identified essential regulators of RPE regeneration

For phenotypic analysis, nitroreductase/metronidazole (NTR/MTZ)-mediated ablation of the RPE was conducted on F0 knockout larvae and scrambled controls carrying the *rpe65a*:nfsB-eGFP transgene (Fig. 1D). RPE markers (e.g., ZPR2) and pigment are largely recovered by 4dpi in control larvae (Hanovice *et al*., 2019; Leach *et al*., 2021; Lu *et al*., 2022), and here, we used these readouts to assess the impact of GOI knockout on RPE regeneration at 4dpi. To facilitate large-scale phenotyping, quantification of normalized brightfield images was performed using RpEGEN, a MATLAB-based platform developed to automate quantification of RPE/pigment recovery after ablation (Leach *et al*., 2022). RpEGEN quantifies 8-bit pixel intensities within an RPE region of interest (ROI), which extends 0 to 180 angular degrees (dorsal to ventral in the eye), and statistically compares results between GOI and scrambled controls. Statistical comparisons spanning 0-30 angular degrees (distal-most dorsal RPE) and 150-180 angular degrees (distal-most ventral RPE) were omitted, as the *rpe65a*:nfsB-eGFP transgene expression and MTZ-induced ablation are confined to the central two-thirds of the RPE (Hanovice *et al*., 2019). To identify phenotypes, we established a selection criterion in which continuous blocks of statistically significant pigment differences (p-value ≤ 0.05) spanning more than twenty angular degrees would be considered phenotypically relevant. Using this criterion, eight GOIs *(cldn7b, fosl1b, nrg1, ogflr1, ccn1l1, lipib, cidec*, and *il11a)* showed angular degree block(s) with significantly higher median pixel intensity (indicating lighter pixels/less pigmentation) in F0 knockout larvae, when compared to scrambled controls (Fig. S2, Table S2). These eight GOIs were therefore determined to be positive regulators of RPE regeneration. Seven genes *(cpa4, zgc:153911, adamtsl7, epha2a, dkk1a, lepb*, and *serpine1)* showed angular degree blocks with significantly lower median pixel intensity (indicating darker pixels/more pigmentation) in F0 knockout larvae, when compared to scrambled controls (Fig. S3, Table S2). These seven GOIs were therefore determined to be negative regulators of RPE regeneration. The remaining 12 genes did not satisfy the criterion defined here for a phenotype (Fig. S4). Figure 2 shows a representative example from each group.

**Fig. 2.**
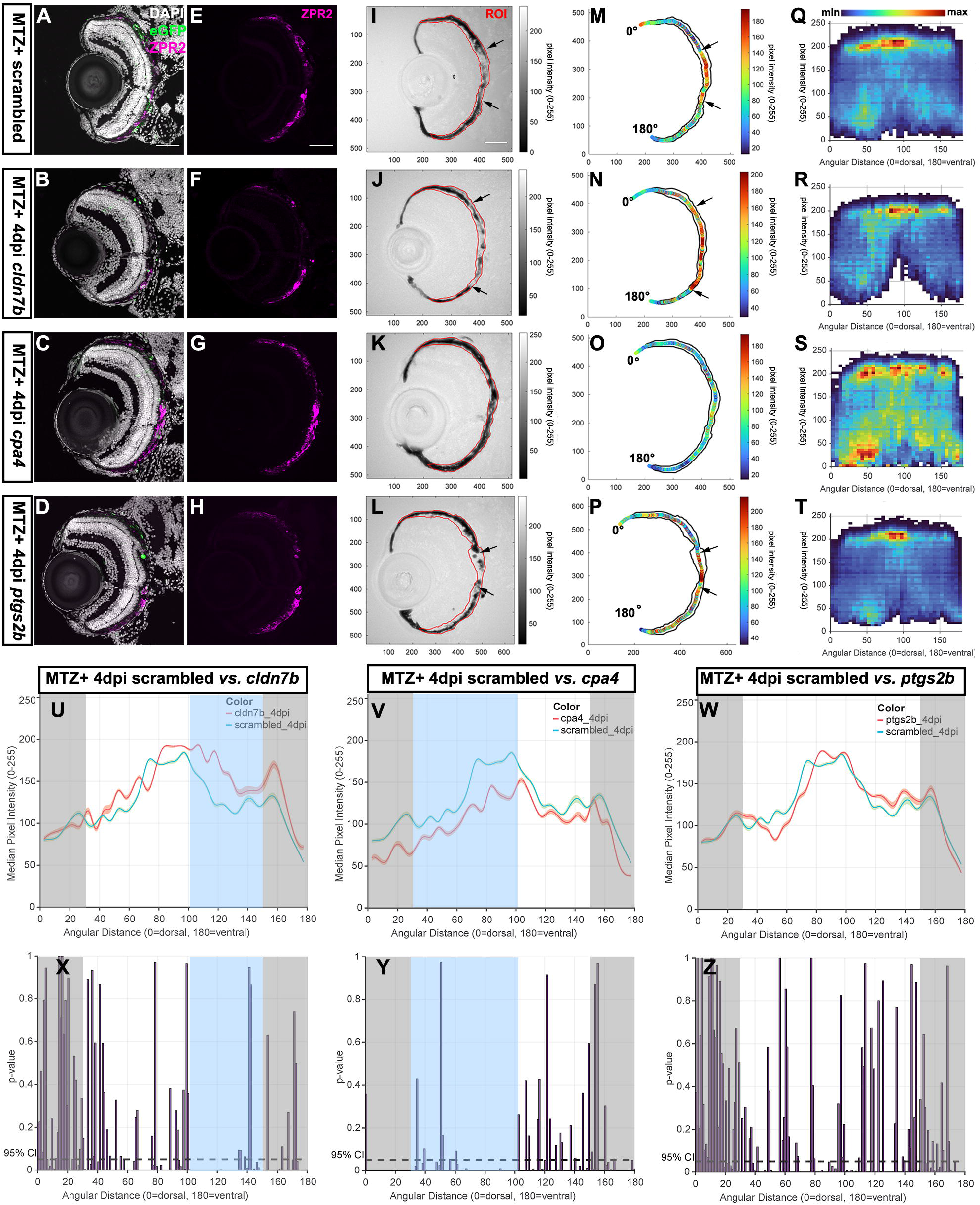
Representative phenotypic screening results by automated quantification of RPE regeneration using RpEGEN. (A-L) Representative immunofluorescence images of ZPR2 staining on transverse cryosections from ablated (MTZ^+^) scrambled, *cldn7b, cpa4*, and *ptgs2b* RNP-injected F0 larvae at 4dpi. Single-channel immunofluorescence images of ZPR2 (E-H) and brightfield (I-L) images. Color bars corresponding to brightfield images indicate an 8-bit scale (0=black, 255=white). Black arrows delineate the edges of pigment recovery. Regions of interest (ROIs) are outlined in red. Nuclei (white), eGFP (green), ZPR2 (magenta). (M-P) Median pixel intensity distributions in the ROI from each group. (Q-T) Heatmaps showing the dorsal-to-ventral (angular distance from 0 to 180°) distribution of raw pixel intensity and (U-W) median intensity pixel values within each ROI from ablated scrambled, *cldn7b, cpa4, and ptgs2b* RNP-injected F0 larvae at 4dpi. Bin size=5 angular degrees. Color bar indicates bin counts. (X-Z) Statistical comparisons of dorsal-to-ventral median pixel intensity between (X) scrambled and *cldn7b* groups, (Y) between scrambled and *cpa4* groups, and (Z) between scrambled and *ptgs2b* groups. Light blue rectangles indicate the region with significant differences spanning more than twenty angular degrees. Gray rectangles indicate the distal-most dorsal (0 to 30 angular degrees) and distal-most ventral (150 to 180 angular degrees) peripheral RPE areas omitted from analyses. Dashed black lines indicate a 95% confidence interval (CI), Bin size=1 angular degree. Scale bar=50 μm. Abbreviations as follows: MTZ, metronidazole; dpi, days post-injury; min, minimum; max, maximum; ROI, region of interest.

While it was determined that 12 GOIs were not essential regulators of RPE regeneration, this assessment was made using pigment recovery as a screening criterion and does not take into consideration other aspects of RPE regeneration such as the recovery of apical/basal polarity or functionality of the regenerated RPE. It is quite possible that using a series of integrated phenotypic screens applied in parallel to the same pool of F0 knockout larvae could further expand this list of regulators of RPE regeneration. F0 screens using pre-screened RNPs have been shown to result in high phenotypic penetrance (>94%) and diversity of null alleles with over 80% of alleles harboring a frameshift mutation (Kroll *et al*., 2021). Here, of the 7 GOIs for which we could not amplify across potential intragenic deletions, 5 showed regeneration phenotypes. Thus, while pre-screening RNPs for those that generate large deletions is helpful in ensuring that all embryos screened possess a mutagenized locus, this is not an absolute requirement for phenotype detection.

The representative positive regulator, *cldn7b*, showed a noticeable expansion of the central RPE injury site, which lacked ZPR2 signal (Fig. 2B,F) and pigmentation (Fig. 2J) in F0 knockout larvae when compared to scrambled controls (Fig. 2A,E,I). Quantitative analysis using RpEGEN showed an extended distribution of lighter pixels with higher raw (Fig. 2R) and median (Fig. 2N) intensity values in the RPE of *cldn7b* F0 knockout larvae when compared to scrambled controls (Fig. 2M,Q). Dorsal-to-ventral RPE pigment quantification revealed significantly higher median pixel intensity from 100 to 150 angular degrees in the *cldn7b* F0 knockout group, when compared to scrambled controls, indicating less pigmentation (Fig. 2U,X). By comparison, the representative negative regulator, *cpa4*, showed both ZPR2 distribution (Fig. 2C,G) and pigment recovery (Fig. 2K) extended further into the central RPE injury site in F0 knockout larvae when compared to scrambled controls (Fig. 2A,E,I). Consistently, *cpa4* F0 knockout larvae also showed an overall distribution of darker pixels with lower intensity values (Fig. 2O,S), and there were significant differences in median pixel intensity from 30 to 106 angular degrees when *cpa4* F0 knockout larvae were compared to scrambled controls (Fig. 2V,Y). Finally, *ptgs2b* F0 knockout larvae, representative of the no phenotype group, showed similar ZPR2 distribution (Fig. 2D,H) and pigment recovery (Fig. 2L) to that of scrambled controls (Fig. 2A,E,I; arrows). Quantification of median pixel intensity values in *ptgs2b* F0 knockout larvae (Fig. 2P,T) did not reveal significant differences with scrambled controls (Fig. 2M,Q,W,Z).

### Macrophages/microglia are retained in the RPE layer of F0 *cldn7b* knockout larvae during regeneration

To further validate the robustness of our F0 screen, we characterized the requirement for *cldn7b* during RPE regeneration. Claudin-7b is a multifunctional protein that maintains apical tight junction (TJ) barrier integrity (Li *et al*., 2018) and epithelial cell polarity (Jin *et al*., 2020) in zebrafish. Claudin-7 also functions in suppressing cell proliferation and migration by modulating cell-matrix interactions (Lu *et al*., 2015; Kim *et al*., 2019; Li and Yang, 2022). Multiple studies using *claudin-7* knockout mice have also highlighted its role in regulating inflammatory responses (Ding *et al*., 2012, 2022; Tanaka *et al*., 2015). As proliferation is a critical post-RPE ablation response (Hanovice *et al*., 2019), we first explored whether knockout of *cldn7b* impacted cell proliferation in regenerating RPE. At 4dpi, the number of proliferating (BrdU-labeled) cells in the RPE layer was significantly increased in the ablated (MTZ+) scrambled and *cldn7b* knockout larvae when compared to corresponding unablated (MTZ-) controls, as expected (Hanovice *et al*., 2019); however, there was no significant difference in proliferating cells between *cldn7b* F0 knockouts and scrambled controls (Fig. S5) indicating cldn7b does not contribute to the proliferative response post-injury.

Innate immunity and inflammation play a critical role during RPE regeneration (Leach *et al*., 2021). In response to injury, macrophages/microglia accumulate in and around the RPE between 2-3dpi, likely acting as scavengers to phagocytose apoptotic bodies and cell debris, and triggering an inflammatory response. Between 3-4dpi, macrophage/microglia accumulation wanes and inflammation likely resolves to facilitate recovery of pigment/RPE (Leach *et al*., 2021). With this in mind, we next sought to investigate whether *cldn7b* knockout affects RPE regeneration by modulating injury-induced immune responses. To test this hypothesis, we utilized *mpeg1*:mCherry transgenics (Ellett *et al*., 2011), to assess macrophage/microglia localization from 2-4dpi. At 2dpi, macrophages/microglia were accumulated in the RPE injury site, but there was no significant difference between ablated *cldn7b* F0 knockout larvae and scrambled controls (Fig. 3A,B,G). By 3dpi, substantial infiltration of mCherry+ macrophages/microglia was observed in the RPE layer in both *cldn7b* F0 knockout larvae and scrambled controls (Fig. 3C,D), as anticipated (Leach *et al*., 2021), and quantification showed there were significantly more macrophages/microglia in the ablated *cldn7b* F0 knockout larvae relative to scrambled controls (Fig. 3G). Interestingly, overlaying the mCherry and brightfield channels revealed a close association between mCherry+ cells and pigment debris (Fig. 3C’’,D’’), which was especially evident in the *cldn7b* F0 knockout larvae where the mCherry+ signal appeared to encompass pigment clusters (Fig. 3D’’). By 4dpi, macrophage/microglia presence started to wane in the scrambled controls (Fig. 3E,H,G), while significant retention of mCherry+ signal was observed in the RPE injury site of *cldn7b* F0 knockout larvae (Fig. 3F,I,G). Notably, there were no significant differences in mCherry signal quantification between the unablated groups at any time point (Fig. S6G-I), indicating a role for *cldn7b* in regulating leukocyte infiltration in response to injury. These observations are consistent with those in *Cldn7* knockout mice where loss of Cldn7 promoted leukocyte infiltration and intestinal inflammation by damaging the intestinal epithelium (Wang *et al*., 2021), increasing intestinal epithelial permeability, and disrupting cell-matrix interactions (Ding *et al*., 2012, 2022; Tanaka *et al*., 2015). The structure of the RPE layer appeared overtly normal in unablated *cldn7b* knockout larvae (Fig, S6), and therefore the increased accumulation of macrophages/microglia in the RPE layer after injury might be related to the non-TJ function of claudin-7 in modulating cell-matrix adhesion.

**Fig. 3.**
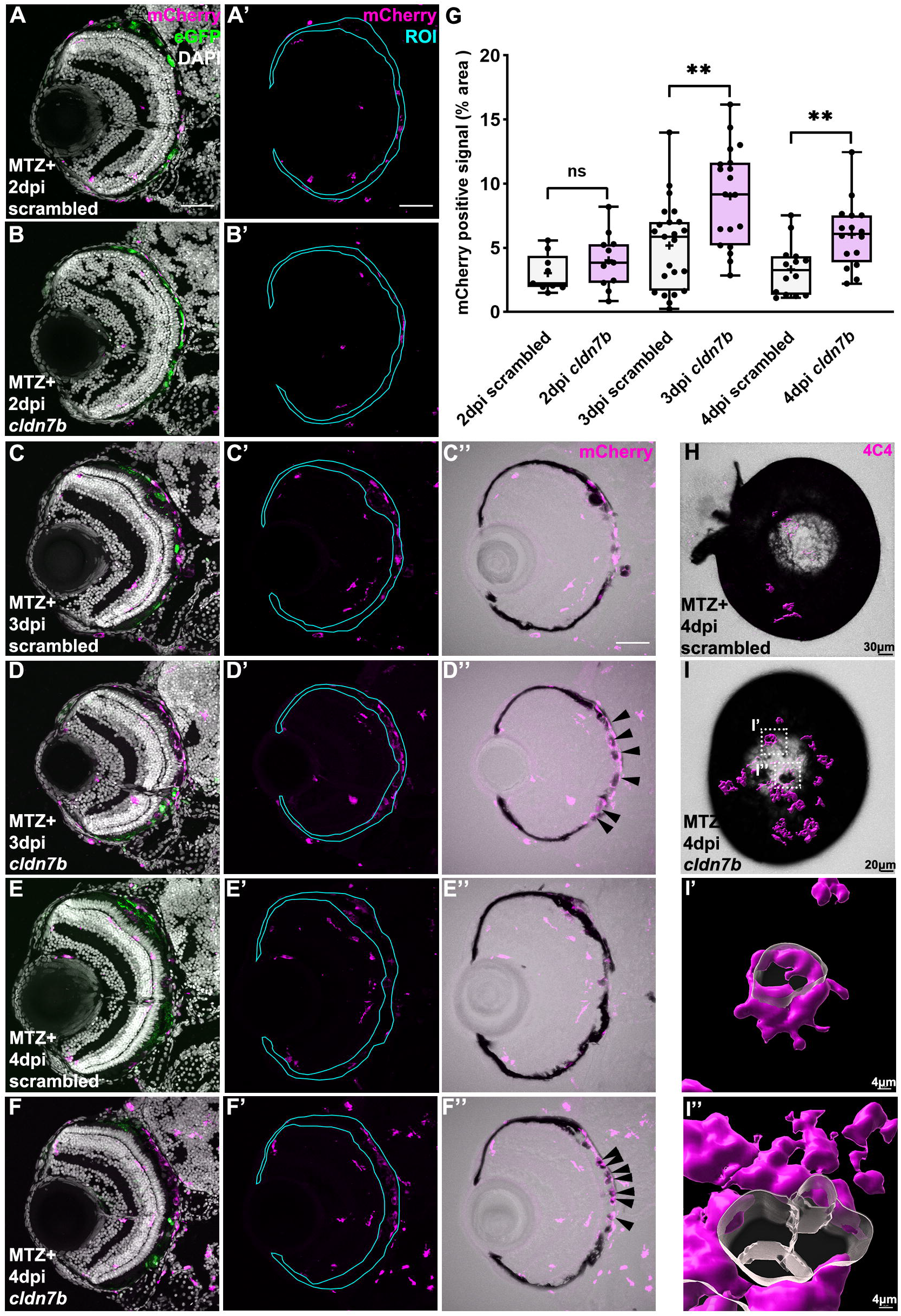
*cldn7b* F0 knockout larvae show retention of macrophages/microglia and impaired clearance of pigment debris during RPE regeneration. (A-F) Representative immunofluorescence images of mCherry signal in ablated (MTZ+) scrambled and *cldn7b* F0 knockout larvae at (A-B) 2dpi, (C-D) 3dpi, and (E-F) 4dpi. (A’-F’) Single-channel images of mCherry signal in the RPE ROIs. (C’’-F’’) Overlaid images of mCherry and bright-field channels. Nuclei (white), eGFP (green), mCherry (magenta). (G) Box plots showing significantly more mCherry+ macrophage/microglia signal in the RPE layer of *cldn7b* F0 knockout larvae at 3dpi and 4dpi, compared to the scrambled controls. (H-I) Overlaid images of 3D-rendered 4C4+ isosurfaces (magenta) with bright-field channels from ablated scrambled and *cldn7b* knockout whole-mount eyes. 4C4+ macrophages/microglia (magenta) surfaces were 3D-rendered in Imaris. (I’, I’’) Zoomed-in views of (I; white boxes) *cldn7b* knockout whole-mount eye showing 4C4+ cells engulfing the pigment debris. (A-F) scale bar=50μm. (H-I) Scale bars were indicated on the images. **P value ≤ 0.01; ns=not significant. Abbreviations as follows: MTZ, metronidazole; dpi, days post-injury; ROI, region of interest.

Pigment was largely recovered in the control larvae at 4dpi (Fig. 3E’’), as anticipated (Hanovice *et al*., 2019), whereas *cldn7b* F0 knockout larvae showed less pigment recovery and the remaining pigment debris appeared to be surrounded by macrophages/microglia retained in the RPE injury site (Fig. 3F’’). Engulfment of pigment debris by macrophages/microglia was further examined using 3D-rendering of whole-mount eyes labeled with 4C4, a marker for zebrafish macrophages/microglia (Craig *et al*., 2008; Rovira *et al*., 2022). Consistent with quantification of sectioned tissue (Fig 3G), these data showed an accumulation of 4C4+ cells in *cldn7b* F0 knockout eyes (Fig. 3I compared to 3H). Moreover, the 4C4+ signal surrounded pigment debris (Fig. 3I-I”), suggesting a possible role for claudin-7b in assisting macrophage/microglia with phagocytosis/removal of degenerating RPE post-ablation.

The expression of claudins is species-specific, tissue-specific, and cell-specific within a tissue, and expression also varies with the stage of development. Claudin expression/distribution is not known in zebrafish, but in cultured human fetal RPE (hfRPE), claudin-19 and claudin-3 are enriched (Peng *et al*., 2011), while in murine RPE, *Cldn1, Cldn2, Cldn4, Cldn7*, and *Cldn23* are all expressed (Louer *et al*., 2020). Macrophages also express claudin-1, claudin-2, claudin-11 (Van den Bossche *et al*., 2012), and claudin-7 (Zheng *et al*., 2005). Claudin-1 in macrophages is predicted to interact with the airway epithelium and help preserve tissue integrity (Blank *et al*., 2011). Claudin-1 is also thought to aid pathogen clearance by macrophages in an *in vitro* intestinal epithelium model (Bording-Jorgensen *et al*., 2021). Phagocytosis of apoptotic cells by macrophages facilitates the expression of anti-inflammatory markers and contributes to the switch to an anti-inflammatory phenotype (Fadok *et al*., 1998; Freire-de-Lima *et al*., 2006). Thus, it is possible that cldn7b deficiency in zebrafish impairs the resolution of inflammation during the later phases of RPE regeneration and thereby impedes regenerative processes. Importantly, our *cldn7b* F0 knockout model cannot distinguish whether cldn7b is required in RPE cells, macrophages/microglia, and/or other cells (like RPE stem cells) (Xing *et al*., 2020) that might be involved in regeneration. Future studies are needed to understand the mechanism by which *cldn7b* regulates RPE regeneration, and these can be performed in stable *cldn7b* mutant lines.

In summary, we report an effective and rapid F0 screening platform to identify regulators of RPE regeneration in zebrafish. From a pilot screen, we identified 15 novel RPE regeneration regulators; further studies of these GOIs will greatly expand our understanding of the regenerative process and some of the candidates might be useful foundations around which therapeutics could be developed to slow or even treat RPE degenerative diseases. Moreover, the screen can be performed at a larger scale to expand even further the number of putative regulators of RPE regeneration.

## MATERIALS AND METHODS

### Ethics statement

All animal experiments were performed with permission from the University of Pittsburgh School of Medicine Institutional Animal Care and Use Committee.

### Zebrafish husbandry

Adult zebrafish *(Danio rerio)* were maintained under standard conditions at 28.5°C in circulating system water on a 14-hour light/10-hour dark cycle. Embryos and larvae (≤ 9dpf) used for subsequent experiments were kept in the dark in an incubator at 28.5°C until being euthanized by tricaine overdose (MS-222; Western Chemical Inc.).

### RPE ablation

RPE ablation was performed using the transgenic *rpe65a:*nfsB-eGFP zebrafish line (mw86Tg) as described using an established protocol (Leach, Fisher and Gross, 2022) Briefly, embryos derived from wide-type AB and *rpe65a:*nfsB-eGFP outcrosses or *mpeg1:*mCherry (gl23Tg) (Ellett *et al*., 2011) and *rpe65a:*nfsB-eGFP outcrosses were treated with 0.003% 1-phenyl 2-thiourea (PTU; Sigma-Aldrich) from 6hpf to 5dpf to prevent pigmentation. At 5dpf, larvae were screened for the *rpe65a:*nfsB-eGFP and/or *mpeg1:*mCherry transgenes and immersed in 10 mM metronidazole (MTZ; Sigma-Aldrich) dissolved in system water for RPE ablation. MTZ was removed after 24-hour treatment and fresh system water was replenished daily until larvae were euthanized for experiments.

### BrdU incorporation assay

For bromodeoxyuridine (BrdU) incorporation experiments, larvae were exposed to system water containing 10 mM BrdU (Sigma-Aldrich) for 24 hours before euthanasia.

### crRNA selection/design and RNP preparation and injection

crRNAs for each target gene were either selected from the Alt-R Predesigned Cas9 crRNA Selection Tool or designed using the Alt-R Custom Cas9 crRNA Design Tool within the Integrated DNA Technologies (IDT) database. crRNAs were selected based on published criteria (Kroll *et al*., 2021) and designed to target three distinct and asymmetric exons shared by all/most transcripts for the target genes. RNPs were prepared as described (Kroll *et al*., 2021); briefly, the crRNA was annealed with an equal molar amount of tracrRNA (IDT, #1072532) in duplex buffer (IDT, #11010301) to form gRNA by heating at 95°C for 5 min and subsequently cooling on ice. gRNA was assembled with an equal molar amount of Alt-R S.p. Cas9 Nuclease V3 (IDT, #1081058) to form the RNP complex (28.5 µM final concentration) by incubation at 37°C for 5 min followed by storage at -20°C. For pre-screening validation of individual RNPs, approximately 1 nl of the single RNP complex (28.5 µM final concentration) was injected into the yolk of one-cell stage embryos before cell inflation. For phenotypic screening, three RNPs (9.5 µM each individual RNP) were pooled to yield a final concentration of 28.5 µM RNPs (or 4700 pg Cas9 and 1000 pg each gRNA per embryo). Approximately 1 nl of the three-RNP complex pool was injected into one-cell stage embryos. To account for any non-specific effects resulting from the injection itself, pooled RNP complexes were prepared from three scrambled crRNAs (IDT: Alt-R CRISPR-Cas9 Negative Control crRNA #1, 1072544; #2, 1072545; #3, 1072546) and injected as described above. Additional details about the design method, target locus, and sequence of each crRNA can be found in Table S1.

### Standard, headloop, and genotyping PCR

Genomic DNA was extracted from embryos or larval tails by incubating with 50ul of 50mM NaOH at 95°C for 20 minutes. After cooling to 4°C, 5ul of 1M Tris-HCl (pH 8) was added for neutralization. For standard PCR, each 25µl reaction contained 1µl of genomic DNA, 1µl 10 µM forward and reverse primer mix, 0.5µl 10 mM dNTPs, 5µl standard 5X Phusion HF Buffer, and 0.1µl of Phusion DNA polymerase (NEB). For headloop PCR, one of the primers was replaced by a 2.5 µM primer tagged with 20 bases that are completely complementary to the target locus. Standard and headloop PCR products from n=10 RNP-injected embryos and n=2 uninjected embryos were run in parallel on a 1% agarose gel for data analysis. For genotyping PCR, the same reaction as standard PCR was used to amplify a short (<1kb) sequence spanning all three target sites. For amplicons >1kb, 25ul reactions were modified to include: 2µl of genomic DNA, 2.5µl 10 µM primer mix, 0.5µl 10 mM dNTPs, 5µl standard 5X Phusion HF Buffer, and 0.2µl of Phusion DNA polymerase. Longer amplicons were analyzed on a 2% agarose gel. Additional details about primers used in standard, headloop, and genotyping PCR can be found in Table S1.

### Quantitative real-time PCR (qRT-PCR)

Total RNA was extracted and purified from pooled scrambled control and genotyped *cldn7b* F0 knockout larvae (n=50 per group) at 7dpf using the RNeasy Plus Micro Kit (QIAGEN), followed by cDNA synthesis with the SuperScript IV VILO Master Mix Kit (ThermoFisher). Forward (5’-AATCCTCTCTGTTGGAGCCCT-3’) and reverse (5’-TTGACAGGTGTGAAGGGGTTG-3’) primers spanning the *cldn7b* exon 2-exon 3 junction were designed using Primer BLAST (https://www.ncbi.nlm.nih.gov/tools/primer-blast/). Each qRT-PCR reaction (10 µl/well) was assembled by adding 2 µl 0.5ng/µl cDNA, 0.5 µl 10 µM forward and reverse primer mix, and 5 µl 2X iTaq Universal SYBR Green Supermix (Bio-Rad Laboratories). Experiments were run in three technical replicates on a CFX384 Touch Real-Time PCR Detection System (Bio-Rad Laboratories). Relative gene expression fold change was calculated using the Livak method (2^− ΔΔCt^) (Livak and Schmittgen, 2001). GAPDH was the housekeeping gene chosen for expression normalization (Barber *et al*., 2005) as *cldn7b* knockout did not affect baseline expression.

### Immunohistochemistry on cryosections and whole-mount larvae

Larvae were fixed in 4% paraformaldehyde (PFA) at room temperature for 2–3 hours or overnight at 4°C. Immunostaining on cryosections was performed following an established protocol (Uribe and Gross, 2007). Briefly, fixed larvae were rinsed in phosphate-buffered saline (PBS) twice, followed by cryoprotection with sucrose (25% to 35% gradient). Subsequently, tissues were embedded in optimal cutting temperature (OCT) compound and rapidly frozen on dry ice. Transverse cryosections were cut at 12-micron thickness and mounted onto poly-L-lysine pre-coated glass slides (Superfrost Plus; Thermo Fisher). For mCherry and ZPR2 staining, slides were rehydrated in PBS and washed three times with PBTD (1X PBS with 0.1% Tween-20 and 1% DMSO). For BrdU staining, an antigen retrieval step, which consisted of incubating sections in 4N HCl for 8 minutes at 37°C, was added before washes in PBTD. Then, sections were blocked for ≥ 2 hours in 5% normal goat serum (NGS) in PBTD (e.g. blocking buffer) before overnight incubation at 4°C with primary antibody diluted in blocking buffer. Sections were subsequently washed three times with PBTD and incubated for 2–3 hours with secondary antibody diluted in blocking buffer. Slides were then washed in PBTD for 3×10 min and 1mg/ml DAPI (1:250, Sigma-Aldrich) was added and incubated for 10 minutes between the first and second washes. Slides were mounted with Vectashield mounting media (Vector Laboratories) and sealed with nail polish.

Whole-mount staining was performed as described (Maves *et al*., 2002). Briefly, fixed tissues were rinsed with PBST (PBS with 0.1% Tween-20) and distilled water, followed by permeabilization with 100% acetone at −20°C for 10 minutes and subsequent rinsing with distilled water and PBST. Tissues were then digested in a collagenase solution (1mg/mL in PBST) for 30 minutes and followed by proteinase K (2mg/mL in PBST) exposure for another 30 minutes before blocking for ≥1 hour blocking in 2% NGS in PBDTX (PBS with 1% bovine serum albumin, 1% DMSO, and 0.1% Triton X-100, pH =7.3). Larvae were then incubated overnight at 4°C with primary antibody diluted in blocking buffer. The next day, larvae were washed withPBDTX four times and incubated for 4 hours at room temperature in secondary antibody diluted in blocking buffer. Nuclei were counterstained with DAPI, which was added for the last 2 hours of the 4-hour secondary incubation. Subsequently, larvae were rinsed three times in PBS. Eyes were dissected using sharpened tungsten wire and mounted onto slides with 0.5% low melting agarose immediately before imaging.

Primary antibodies used in this study include: mouse anti-zpr-2 (1:250, Zebrafish International Resource Center (ZIRC)), mouse anti-mCherry (1:200, Takara Bio/Clontech Laboratories, 632543), rat anti-BrdU (1:250, Abcam, ab6326), and mouse anti-4C4 (1:200, a kind gift of Dr. Peter Hitchcock, University of Michigan School of Medicine, USA). Secondary antibodies used in this study include goat anti-mouse Cy3 (1:250, Jackson ImmunoResearch Laboratories, 115-165-166) and goat anti-rat Cy3 (1:250, Jackson ImmunoResearch Laboratories, 112-165-003). All fluorescent images were captured using an Olympus Fluoview FV1200 laser scanning confocal microscope (Olympus Corporation). Cryosection images were taken under a 40x objective and whole-mount images were taken under a 20x objective with 1.7 zoom (both Olympus Corporation).

### Imaging processing and quantification

Quantification using FIJI (ImageJ (Schindelin *et al*., 2012)): Raw confocal z-stack images (1µm z-step size) were processed for image presentation and quantification as described (Lu *et al*., 2022). To quantify BrdU, z-stack images were maximum-projected and the number of BrdU+ cells in the RPE layer were counted manually using the Cell Counter Plugin. Each data point represents the average number of BrdU+ cells from three consecutive central sections per larva. To quantify mCherry signal, maximum-projected images were first converted into 8-bit TIF images. Using the Polygon Selection Tool, RPE regions of interest (ROIs) were drawn to encompass the space between the outer edge of the photoreceptor outer segments (e.g. the apical RPE/photoreceptor interface) and the basal-most edges of RPE pigment. Thresholding on the mCherry channel was then performed to measure the percent area of mCherry+ signal within the ROI. The same thresholding parameters (40/255, 8-bit scale) were applied to all images to standardize quantification. Each data point represents the quantification of a single, central-most section from each larva.

Quantification using MATLAB: Raw confocal z-stack images (1µm z-step size) were obtained from n=16 scrambled larvae from 3 independent injections and at least n=8 larvae from genotyped F0 GOI knockout larvae. Images were preprocessed into 8-bit grayscale TIFs and RPE ROIs were drawn using FIJI, as described in detail (Leach *et al*., 2022). Image normalization was achieved by first averaging the mean gray values of each brightfield image in the whole dataset (e.g. all scramble control and GOI images), which averaged to 196 (0-255 8-bit scale). Then, the Brightness Tool (FIJI) was used to adjust the mean gray value of each individual brightfield image to 196/255. Subsequently, normalized TIF images and ROI files were processed in MATLAB (R2021b (version 9.11.0) using the RpEGEN scripts, as described in detail (Leach *et al*., 2022). Robust statistical comparisons between scrambled controls and F0 knockout groups were validated and achieved using 20,000 permutation simulations.

### Statistics

All statistical analyses were performed using Prism 9.0 (GraphPad). For box plots, the line and plus within the box represent the median and mean, respectively; the top and bottom whiskers represent the maximum and minimum values, respectively; and each dot represents a biological replicate (one eye from one larva). For statistical tests, D’Agostino-Pearson omnibus normality test was first performed to determine whether data obeyed normal (Gaussian) distributions. For comparisons between two groups, unpaired Student’s t-tests with Welch’s correction were performed on datasets with normal distribution and non-parametric Mann–Whitney tests were performed on datasets that were not normally distributed. For multiple comparisons, Kruskal–Wallis ANOVA followed by Dunn’s multiple comparisons tests were used to determine the significance between groups. Exact p-values, numbers of independent experiments (N), and numbers of biological replicates (n) for each dataset can be found in Table S3.

## ACKNOWLEDGEMENTS

We are grateful to Dr. G. Burch Fisher for technical assistance in MATLAB, to Florence Welch and Isabella Summers for technical support, and to Dr. Hugh Hammer for zebrafish husbandry.

## DATA AVAILABILITY

RNA-seq datasets used for this study are publicly available through NCBI GEO: Series GSE174538 *(Lu et al*., 2022) and Series GSE155294 (Leach *et al*., 2021). The RpEGEN scripts used for quantification in this study are freely available and can be downloaded from the GitHub repository: https://github.com/burchfisher/RpEGEN (Leach *et al*., 2022).

